# BRET-Based Ligand Binding at Recombinant and Endogenous Dopamine D2 Receptors

**DOI:** 10.64898/2026.10.23.761207

**Authors:** Kole Martin, Rajani Nimma, Takudzwa Mabvuta, Sandra Ortigueira, Eddy Sotelo, Maria Majellaro, Prashant Donthamsetti

## Abstract

Dopamine D2 receptor (D2R) regulates diverse biological processes and is an important therapeutic target, underscoring the need for robust methods to quantify ligand binding at this receptor. Bioluminescence resonance energy transfer (BRET)-based ligand binding assays have emerged as a powerful platform for quantifying binding in living cells, but their application to D2R has been limited by the availability of suitable fluorescent tracers. Here, we developed three far-red fluorescent antagonists (CELT-240, CELT-383, and CELT-385) as BRET tracers for D2R. All three compounds produced specific, saturable D2R binding with nanomolar affinity. Further characterization of CELT-385 demonstrated its utility for kinetic and competition binding measurements and revealed high affinity binding at the closely related dopamine D3 receptor (D3R). We also extended BRET-based ligand binding beyond genetically-modified receptors using bystander BRET, enabling measurements at unmodified recombinant and endogenously expressed D2R. Together, these findings provide new tools for quantitative D2R/D3R ligand binding and a template for extending BRET-based ligand binding to endogenously expressed GPCRs.

## Introduction

The neuromodulator dopamine regulates diverse physiological and behavioral processes (movement, learning, cognition)^1^. Dopamine signaling is mediated by five G protein-coupled receptors (GPCRs), which are divided into the D1-like receptors (D1R, D5R) and D2-like receptors (D2R, D3R, D4R)^1^. D2R and its close homolog D3R are important therapeutic targets for various neurological and neuropsychiatric disorders (schizophrenia, Parkinson’s disease, addiction), motivating continued development of ligands with improved pharmacological properties^2^.

Key to D2R ligand development is the ability to directly and quantitatively measure receptor-ligand interactions. Ligand binding has traditionally been measured using radioligands, which provide sensitive and quantitative determination of receptor affinity but require the use and disposal of radioactive materials^3^. Fluorescent ligands offer a non-radioactive alternative, but like radioligands, measurements can be complicated by background from free and nonspecifically associated tracer^4, 5^. Filtration and/or washing are therefore required to remove background tracer and isolate the receptor-bound fraction, which can increase assay complexity, limit throughput, disrupt binding equilibrium, and preclude continuous measurement of ligand association and dissociation kinetics.

BRET-based ligand-binding assays provide an alternative approach for quantifying receptor-ligand interactions^5, 6^. In this system, a luminescent donor (e.g., Nluc) is fused to the extracellular N-terminus of the receptor. Binding of a fluorescent acceptor ligand to the receptor positions the fluorophore in close proximity to the donor protein, resulting in increased BRET. Because BRET is generated upon binding of the fluorescent ligand to the receptor, measurements can be performed in the presence of free tracer without filtration or washing. This approach has been successfully applied to diverse GPCRs^7-9^, demonstrating its broad utility.

Despite their advantages, the application of BRET-based ligand binding to D2R presents several challenges. Prior BRET-based ligand binding studies at D2R have employed fluorescent agonist tracers^10^, which could activate the receptor and promote receptor trafficking, potentially confounding live-cell binding measurements by altering receptor function. Fluorescent antagonist tracers offer an alternative that avoids agonist-induced receptor activation but have yet to be established for BRET-based ligand binding at D2R^5, 6^. In addition, conventional BRET-based ligand-binding assays require genetically engineered, donor-tagged receptors. Eliminating the need for receptor modification preserves the native receptor sequence and creates opportunities to measure ligand binding at endogenously expressed receptors within their native cellular environments.

Here we developed and characterized three fluorescent D2R antagonists, CELT-240, CELT-383, and CELT-385, as tracers for live-cell BRET-based ligand-binding measurements. We established specific, saturable binding at D2R and further characterized CELT-385 through equilibrium, kinetic, functional, and competition binding measurements. We further evaluated CELT-385 across dopamine receptor subtypes and established high affinity binding at D3R. We then extended these tracers to a bystander BRET configuration in which Nluc was localized to the plasma membrane independently of the receptor, enabling ligand binding to be measured without direct receptor modification. This approach was also capable of detecting ligand binding at endogenously expressed D2R/D3R in neuronal-like SH-SY5Y cells. Together, these studies establish fluorescent antagonist tracers for quantitative BRET-based D2R ligand binding and extend this approach from genetically-modified receptors to receptors in their native cellular context.

## Materials and Methods

### Cell lines

Human embryonic kidney 293T (HEK293T) were originally obtained from the American Type Culture Collection (ATCC, CRL-3216) and SH-SY5Y cells were graciously provided by the Niswender lab at Vanderbilt University. HEK293T cells were maintained in Dulbecco’s modified Eagle’s medium (DMEM; high glucose) supplemented with 10% (v/v) fetal bovine serum (FBS) at 37°C in a humidified incubator with 5% CO_2_. SH-SY5Y cells were maintained in 45% DMEM and 45% Ham’s F12 medium supplemented with 10% (v/v) fetal bovine serum (FBS) at 37°C in a humidified incubator with 5% CO_2_. Cells were passaged every 2-3 days using trypsin-EDTA and were used between passages 5 and 40. Cell line identity was confirmed by the supplier. Both cell lines are of human origin. Sex is not specified by the supplier and is not considered biologically relevant for the experiments performed.

### Chemicals

CELT-240, CELT-383, and CELT-385 were synthesized at Celtarys Research SL. The chemical structures of these compounds, including their D2-like receptor antagonist pharmacophore and linkers, are not disclosed due to proprietary considerations. Sulpiride and raclopride (Tocris Bioscience), aripiprazole (MedChemExpress), bifeprunox (Cayman Chemical Company), apomorphine (Sigma Aldrich), and furimazine (Selleck Chemicals) were obtained commercially.

### Molecular Biology and Heterologous Expression

All constructs were cloned into mammalian expression vectors. For the BRET-based assays used in this study, 1.2 million HEK293T cells were seeded in 6-well plates, and 800,000 SH-SY5Y cells were seeded in 12-well plates. These cells were then transfected with Lipofectamine 2000 (4 µL/well for 6-well plates or 2 µL/well for 12-well plates) and then washed and harvested in Dulbecco’s phosphate-buffered saline (DPBS) the following day.

All constructs were cloned into mammalian expression vectors. For luminescence emission, saturation binding, binding kinetics, and competition binding experiments, HEK293T cells were transiently transfected overnight using Lipofectamine 2000 with Nluc-D1R (0.6 µg), Nluc-D2R (0.6 µg), Nluc-D3R (0.6 µg), Nluc-D4R (0.6 µg), or Nluc-D5R. For bystander BRET ligand binding experiments, HEK293T cells were transfected with unmodified D2R (2.1 µg) and membrane-anchored Nluc (0.3 µg), whereas SH-SY5Y cells were transfected with membrane-anchored Nluc (1.4 µg) alone. For BRET-based G protein recruitment experiments, HEK293T cells were transfected with unmodified D2R (2.1 µg), Nluc-miniG_si_ (0.06 µg), and membrane-anchored mVenus (5.25 µg).

### Fluorescence and Luminescence Emission Spectra

For fluorescence emission spectra, CELT-240, CELT-383, or CELT-385 were added to black 96-well microplates (Greiner Bio-One). Fluorescence emission spectra were collected using a CLARIOstar Plus plate reader (BMG Labtech) following excitation at 470 nm. For luminescence emission spectra, HEK293T cells expressing Nluc-D2R were harvested, resuspended in DPBS, and dispensed into white 96-well microplates at 24,000 cells per well. Cells were incubated with CELT-240, CELT-383, or CELT-385 (100 nM) for 1.5 hrs. Furimazine was added to a final concentration of 5 μM 3 min prior to measurement. Luminescence emission spectra were collected using a CLARIOstar Plus plate reader.

### BRET-based Ligand Binding Assays

For BRET ligand-binding assays, transfected HEK293T cells were washed, harvested, and resuspended in DPBS at room temperature. Unless otherwise indicated, cells were dispensed into white 96-well microplates (Greiner Bio-One) at 24,000 cells per well and incubated with fluorescent ligand for 1.5 hrs. Furimazine was added to a final concentration of 5 μM 3 min prior to measurement. Luminescence was measured using a CLARIOstar Plus plate reader (BMG Labtech) at 475 ± 30 nm for Nluc donor emission and 660 ± 100 nm for CELT acceptor emission. BRET was calculated as the ratio of acceptor to donor emission.

For saturation binding experiments, cells were incubated with increasing concentrations of CELT-240, CELT-383, or CELT-385 in the absence or presence of 100 μM raclopride to determine total and nonspecific binding, respectively. Specific binding was calculated by subtracting nonspecific from total binding and analyzed by nonlinear regression using a one-site specific binding model. For kinetic measurements, association of 75 nM CELT-385 with Nluc-D2R was monitored continuously for 1.5 hrs. Following equilibration with 75 nM CELT-385 for 1.5 hrs, dissociation was initiated by addition of 100 μM raclopride. Association and dissociation data were fit by nonlinear regression to determine k_*obs*_ and k_*off*_, respectively. The association rate constant (k_*on*_) was calculated as (k_*obs*_ - k_*off*_)/[CELT-385], and the kinetically derived equilibrium dissociation constant (K_*D*_) was calculated as k_*off*_/k_*on*_. Competition binding was performed using 10 nM CELT-385 and increasing concentrations of bifeprunox, aripiprazole, sulpiride, raclopride, or apomorphine.

For bystander BRET experiments, cells expressing unmodified D2R and membrane-anchored Nluc were incubated with 100 nM CELT-240, CELT-383, or CELT-385. For endogenous receptor binding, SH-SY5Y cells were treated with 100 μM retinoic acid for 24 h to promote cell differentiation and D2R/D3R expression, consistent with previous studies^11^. These cells were then transfected with membrane-anchored Nluc and incubated with 10 nM CELT-385 for 1.5 h before BRET measurement.

For Z′-factor determination, transfected cells were treated with vehicle, 100 nM CELT-385, or 100 nM CELT-385 in the presence of 100 μM raclopride. Each condition was evaluated using 12 technical replicates. Assay robustness was evaluated by calculating the Z′ factor according to:

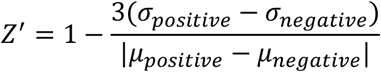

where σ_*positive*_ and σ_*negative*_ represent the standard deviations and *μ*_*positive*_ and *μ*_*negative*_ represent the means of the positive and negative control conditions, respectively^12^.

### BRET-based G protein Recruitment Assay

Consistent with our previous studies^13^, unmodified D2R, Nluc-miniG_si_, and a membrane-anchored mVenus acceptor were co-expressed in HEK293T cells overnight. The following day, cells were detached, resuspended in DPBS, and dispensed into white 96-well plates at 24,000 cells per well. Cells were treated with vehicle, 10 μM dopamine, 100 nM CELT-385, or 100 μM raclopride for 1.5 hrs prior to BRET measurement.

### Data Analysis

All experiments were performed in technical duplicate and independently repeated on at least three separate days using independent cell preparations. Data were analyzed using GraphPad Prism version 11. Nonlinear regression models used for ligand-binding and concentration-response experiments are described above. Comparisons among multiple groups were performed using one-way ANOVA followed by Tukey’s multiple comparisons test. Data are presented as mean ± S.E.M. The number of independent experiments, statistical tests, and significance thresholds are indicated in the corresponding figure legends.

## Results

### Characterization of fluorescent antagonist tracers for BRET-based D2R ligand binding

To identify fluorescent antagonist tracers suitable for BRET-based D2R ligand binding, we developed and characterized three compounds, CELT-240, CELT-383, and CELT-385. Each compound contains the same D2-like receptor antagonist pharmacophore conjugated to the far-red fluorophore BDP-TR through chemical linkers of varying lengths (CELT-383 < CELT-385 < CELT-240). BDP-TR was selected because far-red fluorophores are well suited for BRET-based assays, as their emission is spectrally separated from the blue emission of Nluc, minimizing donor bleed-through and improving signal-to-background^14^. As expected, fluorescence emission spectra confirmed that all three compounds exhibited similar far-red emission profiles (**Supplementary Fig. 1**).

**Figure 1.**
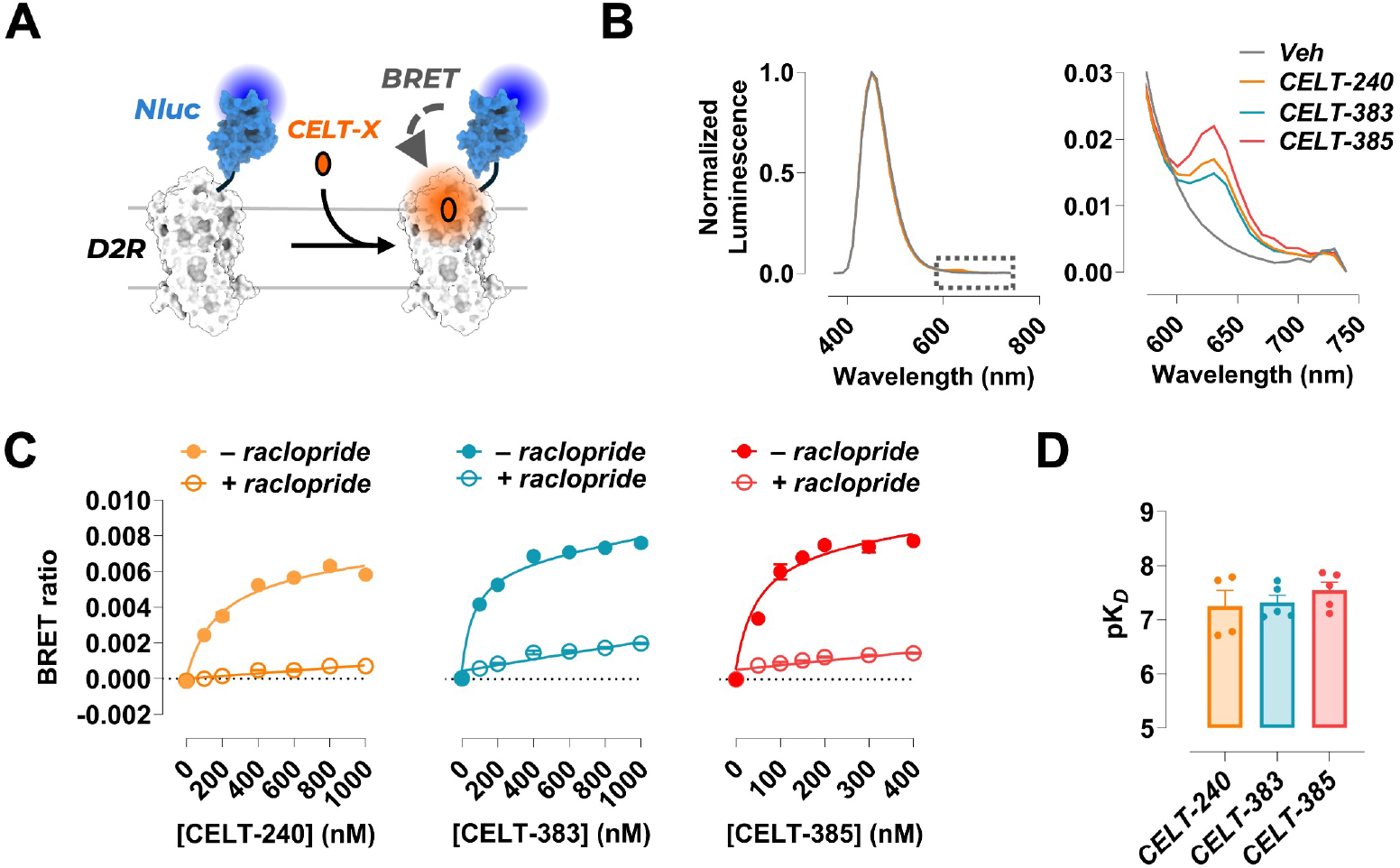
Characterization of a BRET-based binding assay for D2R. (**A**) A BRET-based ligand binding assay between Nluc-tagged D2R (Nluc-D2R) and the fluorescent CELT tracer (direct BRET). (**B**) Luminescence spectra of CELT compounds normalized to peak Nluc emission at 440 nm (*left*). Inset highlights the acceptor emission, which peaks at 630 nm (*right*). (**C**) Representative saturation binding curve of CELT compounds at D2R using the direct BRET-based binding assay. (**D**) Summary of binding affinities (pK_*D*_) of each CELT compound in the direct BRET-based saturation binding assay. one-way ANOVA, Tukey’s post-hoc test, F = 0.67. No significant differences. n = 4 or 5 independent replicates. pK_*D*_ values are as follows: CELT-240 = 7.25, CELT-383 = 7.32, CELT-385 = 7.55.

To determine whether the CELT ligands could function as tracers for live-cell BRET binding, we expressed D2R with Nluc fused to its extracellular N-terminus (Nluc-D2R) in human embryonic kidney 293T (HEK293T) cells, positioning the luminescent donor in proximity to a receptor-bound fluorescent tracer in the orthosteric binding site (**Fig. 1A**). Luminescence emission spectra from Nluc-D2R-expressing cells incubated with each CELT ligand showed the Nluc donor emission peak together with a distinct CELT acceptor emission peak that was absent from donor-only samples (**Fig. 1B**).

CELT-240, CELT-383, and CELT-385 each produced concentration-dependent, saturable BRET signals at D2R that were substantially reduced in the presence of excess the antagonist raclopride (100 μM), demonstrating receptor-specific binding (**Fig. 1C; Supplementary Fig. 2**). All three tracers bound D2R with nanomolar affinity, with pK_*D*_ values of 7.25, 7.32, and 7.55 for CELT-240, CELT-383, and CELT-385, respectively (**Fig. 1D**). Together, these results establish each compound as fluorescent tracers for live-cell BRET binding at D2R.

**Figure 2.**
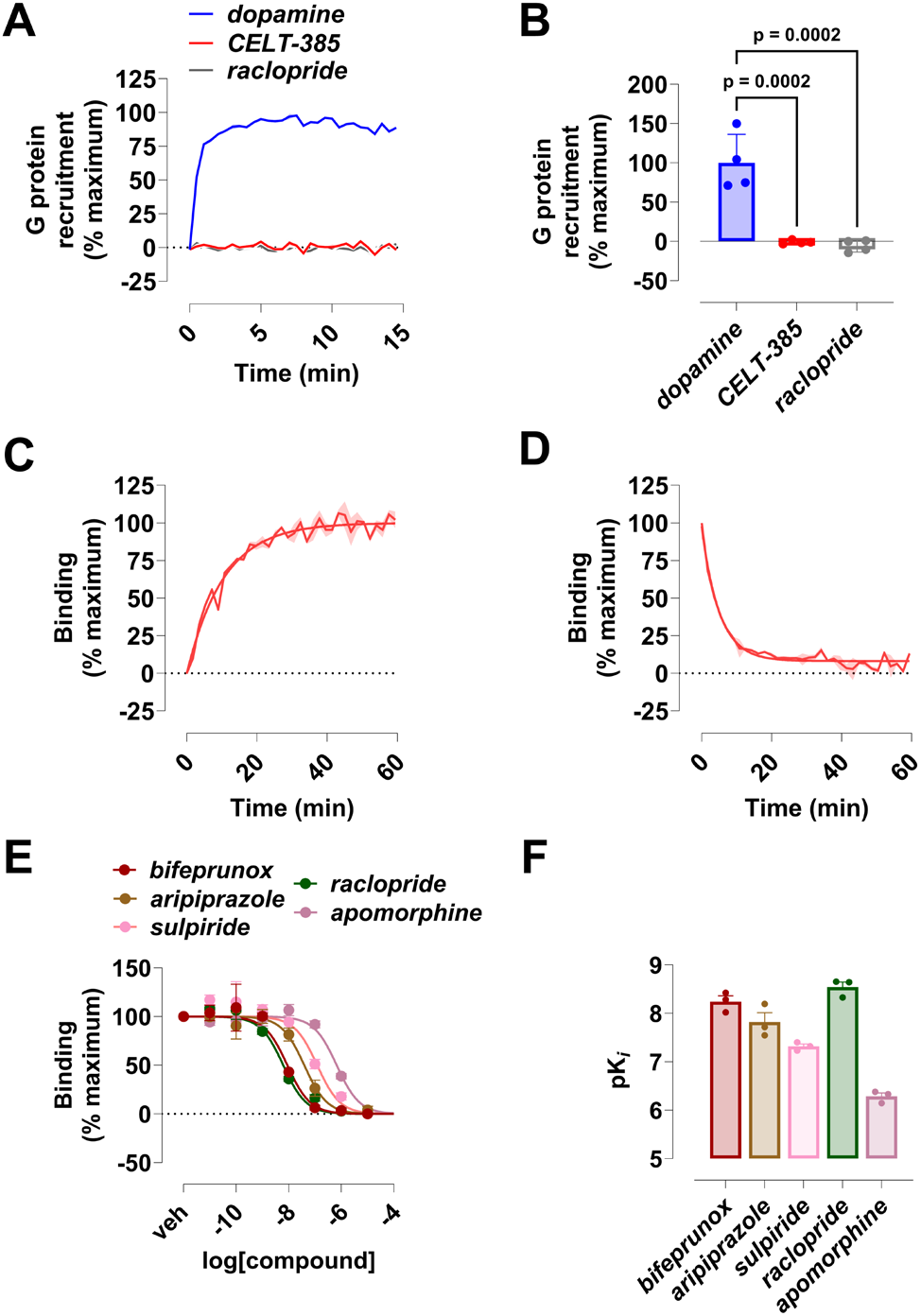
Further characterization of CELT-385. (**A**) Representative G protein recruitment assay at Nluc-D2R. Cells expressing Nluc-D2R were incubated with dopamine (10 µM), CELT-385 (100 nM), or raclopride (100 µM). (**B**) Summary of D2R-mediated G protein recruitment. one-way ANOVA, Tukey’s post-hoc test, F = 31.0. (**C, D**) Representative trace of CELT-385 (75 nM) association (C) or dissociation (D) from Nluc-D2R. An excess of raclopride (100 µM) was used to displace CELT-385 in panel D. n = 4 independent replicates. (**E**) Representative competition binding curves for D2R ligands. Cells expressing Nluc-D2R were incubated with a fixed concentration of CELT-385 (10 nM) and increasing concentrations of each D2R ligand. Binding is expressed as a percentage of maximum fluorescent ligand binding. (**F**) Summary of pK_*i*_ values for D2R ligands. n = 3 independent replicates.

### Extended pharmacological characterization of CELT-385

To further characterize the BRET ligand-binding system, we focused on CELT-385 because it exhibited the highest apparent maximum BRET signal and affinity among the three tracers. We first determined whether CELT-385 behaves as an antagonist at D2R using a G protein recruitment assay^13, 15^. Dopamine induced robust G protein recruitment to D2R, whereas CELT-385 and the antagonist raclopride had no effect at saturating concentrations (**Fig. 2A,B; Supplementary Fig. 3**), consistent with antagonist activity.

**Figure 3.**
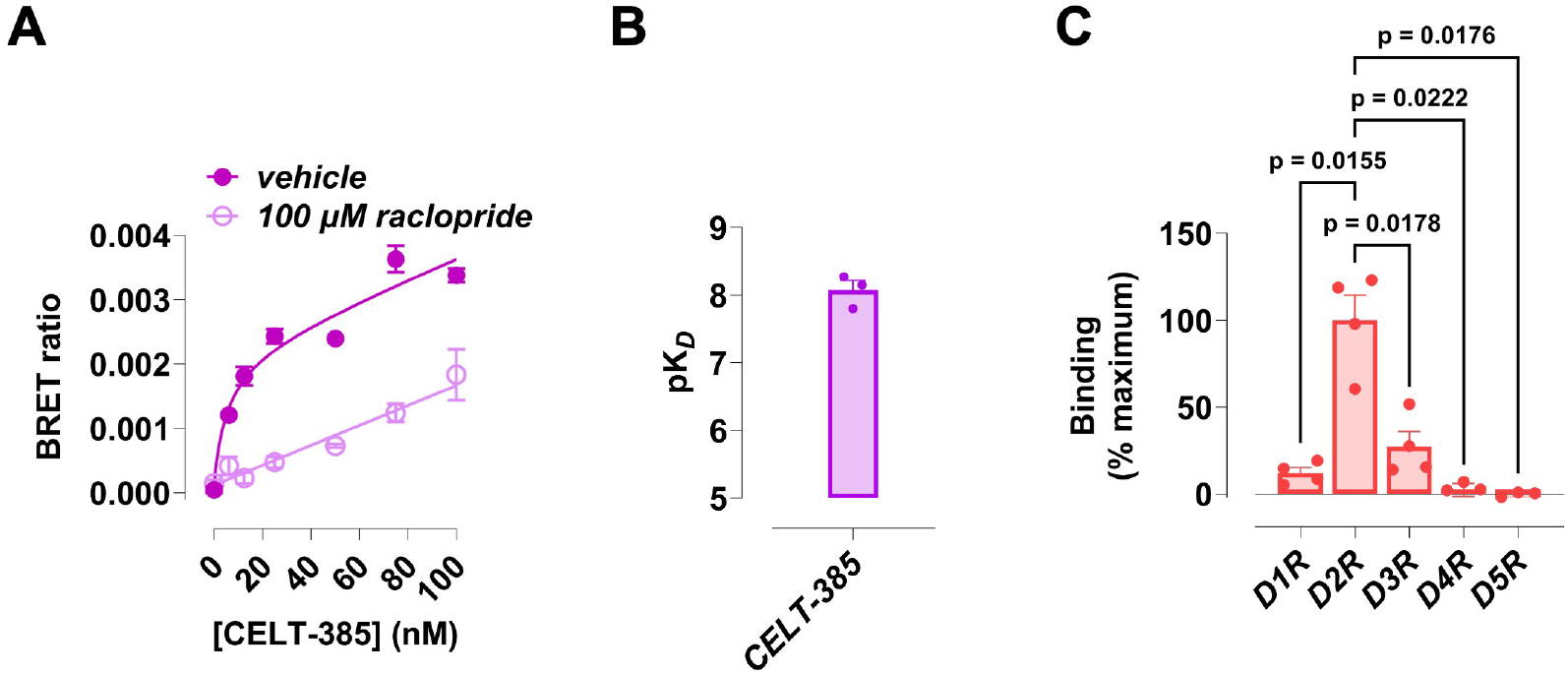
Characterization of CELT-385 binding across dopamine receptor subtypes. (**A**) Representative saturation binding curve of CELT-385 at Nluc-D3R. (**B)** Summary of pK_*D*_ for CELT-385 at Nluc-D3R. n = 3 independent replicates. pK_*D*_ = 8.07. (**C**) Summary of the maximum BRET signal across dopamine receptor subtypes. RM one-way ANOVA, Dunnett’s multiple comparisons test, F = 34.7. n = 4 independent replicates.

We next characterized the binding kinetics of CELT-385 at D2R. Following addition of CELT-385 (75 nM), BRET progressively increased and reached equilibrium (**Fig. 2C**). For dissociation measurements, cells were equilibrated with CELT-385 (75 nM) before addition of excess raclopride (100 μM) to prevent rebinding, resulting in a progressive decrease in BRET as CELT-385 dissociated from D2R (**Fig. 2D**). Analysis of the association and dissociation kinetics yielded a kinetically derived pK_*D*_ (7.35) that was comparable to the affinity determined by equilibrium saturation binding (**Supplementary Table 1**), providing independent support for the measured affinity of CELT-385 at D2R.

We also evaluated the potential of CELT-385 for screening applications. We first assessed assay performance by comparing total and nonspecific binding conditions, which produced a Z′ factor of 0.66 (**Supplementary Fig. 4**), supporting the suitability of CELT-385 for quantitative ligand profiling and higher throughput applications. We then performed competition binding analyses on a panel of unlabeled D2R ligands, including the antagonists raclopride and sulpiride, as well as the partial agonists aripiprazole, bifeprunox, and apomorphine. All five ligands displaced 10 nM CELT-385 from D2R in a concentration-dependent manner, with measured affinities consistent with previously reported values^16-30^ (**Fig. 2D,E; Supplementary Table 2**).

**Figure 4.**
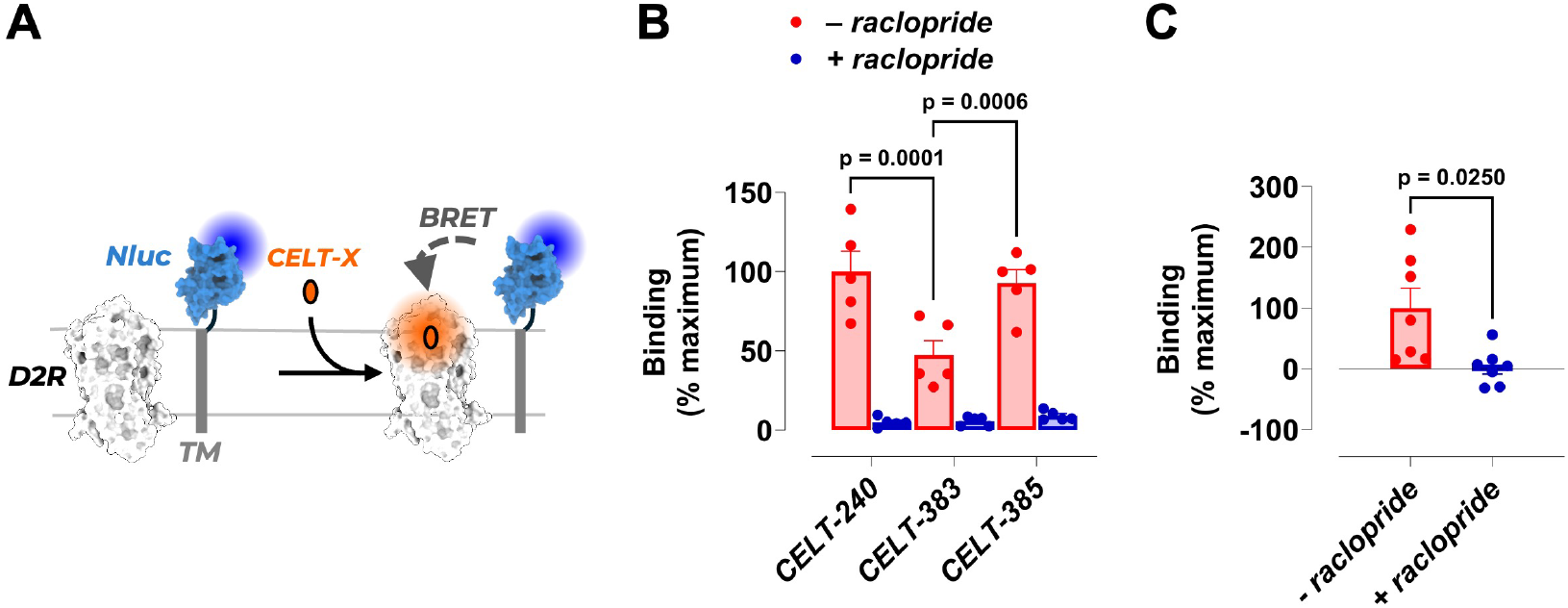
Bystander BRET-based ligand binding at recombinant D2R and endogenous D2R/D3R. (**A**) A BRET-based ligand binding assay between a membrane-anchored Nluc donor that is anchored to the plasma membrane via a single-pass transmembrane segment (Nluc-TM) and a fluorescent CELT tracer that is bound to unmodified D2R. (**B**) Summary of CELT tracer binding to unmodified D2R in HEK293T cells, according to the bystander BRET-based ligand binding assay. two-way ANOVA, Sidak’s multiple comparisons test, n = 5 independent replicates. (**C**) Summary CELT-385 binding to endogenous D2R/D3R in SH-SY5Y neuronal-like cells, according to the bystander BRET-based ligand binding assay. unpaired t-test, n = 5 independent replicates.

Finally, we examined the dopamine receptor subtype binding profile of CELT-385. CELT-385 produced specific, saturable binding at D3R, with a K_*D*_ of ∼8.5 nM (**Fig. 3A-C**) and an ∼2.7-fold preference for D3R over D2R. However, despite its higher affinity at D3R, CELT-385 produced a smaller BRET assay window than at D2R, suggesting a more favorable donor-acceptor geometry at D2R. Across the remaining dopamine receptor subtypes, CELT-385 produced substantially lower specific BRET signals at D1R, D4R, and D5R than at D2R (**Fig. 3A**), indicating preferential recognition of D2R among these dopamine receptor subtypes. Together, these findings establish CELT-385 as a robust fluorescent tracer for quantitative BRET-based pharmacology at D2R and D3R.

### Bystander BRET enables ligand binding at recombinant and endogenous D2R

Conventional BRET ligand-binding assays require fusion of Nluc to the extracellular N-terminus of the receptor, which introduces a substantial modification that could alter receptor function. To determine whether the CELT tracers could detect ligand binding without direct receptor modification, we used a bystander BRET configuration in which Nluc was anchored to the plasma membrane via a single-pass transmembrane segment derived from the low-density lipoprotein receptor (**Fig. 4A**), which we previously showed is preferentially enriched at the plasma membrane^31, 32^. In this configuration, lateral diffusion of the unmodified receptor and membrane-anchored Nluc results in stochastic encounters that transiently position the donor and receptor-bound fluorescent tracer within a distance required for BRET. All three CELT tracers generated receptor-specific bystander BRET at unmodified D2R in HEK293T cells, with CELT-240 and CELT-385 producing significantly greater signals than CELT-383 (**Fig. 4B**). Because Nluc is distributed throughout the plasma membrane, nonspecifically membrane-associated tracer could also come into proximity with the donor and generate BRET. However, BRET measured in the presence of excess raclopride was low, indicating minimal background arising from nonspecific membrane association of the tracers (**Fig. 4B**).

To determine whether bystander BRET could be extended beyond recombinant receptor systems, we next evaluated CELT-385 binding to endogenously expressed D2R/D3R in the neuroblastoma-derived cell line SH-SY5Y. CELT-385 produced measurable BRET signal in SH-SY5Y cells that was significantly abolished in the presence of raclopride (**Fig. 4C**). These findings demonstrate that bystander BRET can detect fluorescent ligand binding at endogenously expressed D2R/D3R in a neuronal-like cellular system.

## Discussion

Here we established fluorescent antagonist tracers for quantitative BRET-based ligand binding at D2R and extended this approach to receptors that are not directly modified with a luminescent donor. Previous BRET-based ligand-binding studies at D2-like receptors have employed fluorescent agonist tracers^10^, which can activate the receptor and alter receptor trafficking during live-cell measurements. The CELT tracers therefore expand the available toolkit by providing antagonist tracers that permit receptor binding to be monitored without simultaneously inducing agonist-dependent receptor activation. Among these, CELT-385 supported equilibrium, kinetic, and competition binding measurements and demonstrated assay performance suitable for compound profiling and higher throughput applications.

CELT-385 also exhibited high-affinity binding at D3R. Interestingly, CELT-385 produced a smaller BRET assay window at D3R despite exhibiting higher affinity for D3R than D2R. This finding highlights an important feature of BRET-based ligand-binding measurements. Signal amplitude depends on factors such as receptor and donor abundance and donor-acceptor distance and orientation. Thus, differences in BRET magnitude between receptor subtypes should be interpreted independently from differences in ligand affinity.

Bystander BRET has previously been applied to fluorescent ligand binding at unmodified GPCRs using membrane-localized *Gaussia* luciferase donor^33^. Our results extend this strategy to Nluc, including ligand binding at endogenously expressed receptors. Notably, bystander BRET signal differed among the tracers despite the use of the same membrane-anchored donor protein. This suggests that bystander BRET signal is influenced not only by receptor occupancy but also by tracer-dependent properties, including linker length, which can affect the position and orientation of the fluorophore relative to the membrane-anchored donor. Tracer selection and fluorophore positioning are therefore important determinants of assay performance in bystander configurations.

Membrane localization of the donor in bystander BRET also creates a potential source of nonspecific signal because membrane-associated fluorescent ligand could be brought into proximity with Nluc. However, the CELT tracers produced little background BRET despite localization of Nluc to the plasma membrane, suggesting minimal contribution from nonspecific membrane association under these conditions. Previous implementations of bystander ligand binding have reported greater contributions from nonspecific signal^33^, suggesting that background may vary considerably depending on tracer properties (linker, fluorophore), donor configuration, and cellular context.

The ability to extend bystander BRET measurements to endogenous receptors is particularly notable because D2R exists within heterogeneous cellular and subcellular environments. In neurons, D2R is distributed across somatic, dendritic, and axonal compartments, where membrane composition, receptor abundance, and local protein interactions can differ^1^. Receptor organization may also influence pharmacology, as previous studies have implicated D2R homo- and heteromerization in ligand binding and receptor function^34-37^. Bystander BRET therefore provides an opportunity to examine ligand binding within native cellular environments, where local membrane composition and receptor organization may influence receptor pharmacology. The affinity of CELT-385 for D3R also highlights the importance of tracer selectivity when extending bystander BRET to endogenous receptor populations, where multiple pharmacologically related receptors may be present.

Decoupling the BRET donor from the receptor also creates opportunities to independently control donor localization. Rather than broadly anchoring Nluc to the plasma membrane, the donor could be targeted to specific membrane proteins using antibodies or other affinity reagents, or directed to defined subcellular compartments. Such approaches could provide spatial control over the receptor populations sampled by bystander BRET and potentially enable ligand binding to be examined within distinct cellular microenvironments. Together, this study expands the fluorescent toolkit for quantitative D2R/D3R ligand binding and provides a framework for extending BRET-based pharmacology to endogenously expressed GPCRs.

## Acknowledgments

This work was supported by Brain and Behavior Research Foundation NARSAD 31429 and NIH grant DP1DA063097-01 (to P.D.). We would like to thank Celtarys for providing the CELT compounds for this study and the Niswender lab at Vanderbilt University for providing the neuronal-like SH-SY5Y cells.

## Author contributions

### Experimentation

K.M. and R.N. Data Analysis: K.M., R.N., and P.D. Chemical synthesis: S.O., E.S., and M.M. Manuscript Preparation: K.M. and P.D. Funding acquisition: P.D.

