## Supplement for "BRET-Based Ligand Binding at Recombinant and Endogenous Dopamine D2 Receptors"

### Supplemental Figures

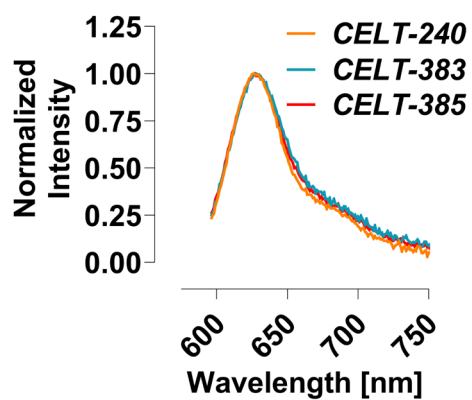

**Supplementary Figure 1. Fluorescence emission spectra of CELT tracers.** Emission spectra of BDP-TR-containing CELT tracers were measured following excitation at 470 nm. Fluorescence intensity was normalized to the maximum emission intensity of each compound (630 nm).

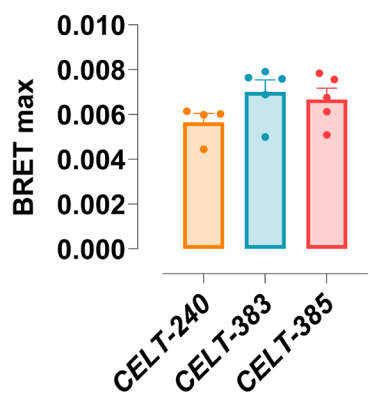

**Supplementary Figure 2. Maximum specific binding in the BRET-based D2R ligand binding assay.** Summary of the maximum specific BRET signal across CELT tracers at Nluc-D2R. one-way ANOVA, Tukey's post-hoc test, $F = 1.9$. No significant differences. $n = 4$ or 5 independent replicates.

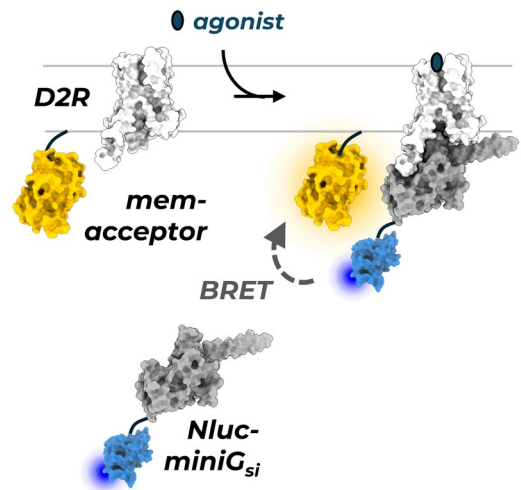

**Supplementary Figure 3. A BRET-based D2R-mediated G protein recruitment assay.** Schematic of a bystander BRET-based G protein recruitment assay. The Gα subunit was engineered to operate independently of G$\beta\gamma$, localize to the cytosol rather than the membrane, and is fused to an Nluc donor (Nluc-miniG<sub>si</sub>). Receptor activation results in the recruitment of Nluc-miniG<sub>si</sub> to D2R, resulting in bystander BRET with a membrane-anchored acceptor (mem-acceptor).

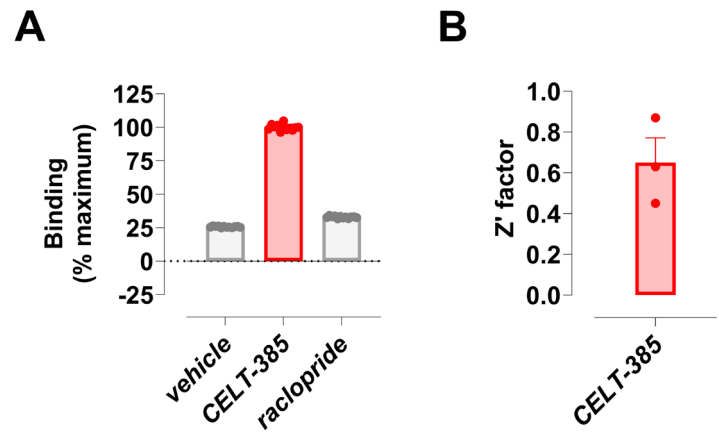

**Supplementary Figure 4. Z' analysis of CELT-385 binding to Nluc-D2R.** (A) Representative binding of CELT-385 (100 nM) to Nluc-D2R in the absence and presence of saturating raclopride (100 μM). n = 12 technical replicates per condition. (B) Summary of Z' factor for CELT-385. n = 3 independent experiments.

### Supplemental Tables

**Supplementary Table 1. Kinetic parameters of CELT-385 for Nluc-D2R.**

| **$K_{on}$ ($10^{-6} \text{ M}^{-1} \text{ min}^{-1}$)** | **$K_{off}$ ($\text{min}^{-1}$)** | **$pK_D$** |
| --- | --- | --- |
| 0.0049 ± 0.003 | 0.22 ± 0.1 | 7.35 |

**Supplementary Table 2. Binding affinities of unlabeled D2R ligands.**

| Compound | pK<sub>i</sub> |  |
| --- | --- | --- |
|  | *This study* | *Literature\** |
| *bifeprunox* | 8.2 ± 0.1 | 8.6 ± 0.5<sup>20, 22, 26</sup> |
| *aripiprazole* | 7.8 ± 0.2 | 8.2 ± 0.2<sup>20, 21, 26</sup> |
| *sulpiride* | 7.3 ± 0.1 | 8.0 ± 0.1<sup>18, 19, 27</sup> |
| *raclopride* | 8.5 ± 0.1 | 8.8 ± 0.1<sup>16, 17, 27</sup> |
| *apomorphine* | 6.3 ± 0.1 | 6.8 ± 0.1<sup>28-30</sup> |

\*Literature pK<sub>i</sub> values are presented as the mean ± S.E.M. of values reported across three independent studies for each compound.
